# Dark diversity reveals importance of biotic resources and competition for plant diversity across broad environmental gradients

**DOI:** 10.1101/685040

**Authors:** Camilla Fløjgaard, Jose W. Valdez, Lars Dalby, Jesper Erenskjold Moeslund, Kevin K. Clausen, Rasmus Ejrnæs, Meelis Pärtel, Ane Kirstine Brunbjerg

## Abstract

Species richness is the most commonly used metric to quantify biodiversity. However, examining dark diversity, the group of missing species which can potentially inhabit a site, can provide a more thorough understanding of the processes influencing observed biodiversity and help evaluate the restoration potential of local habitats. So far, dark diversity has mainly been studied for specific habitats or largescale landscapes while less attention has been given to variation across broad environmental gradients or as a result of local conditions and biotic interactions. In this study, we investigate the importance of local environmental conditions in determining dark diversity and observed richness in plant communities across broad environmental gradients. We use the ecospace concept to investigate how abiotic gradients (defined as position), availability of biotic resources (defined as expansion), spatiotemporal extent of habitats (defined as continuity), as well as species interactions through competition, relate to these biodiversity measures. Position variables were important for both plant richness and dark diversity, some with quadratic relationships, e.g., plant richness showing a unimodal response to soil fertility corresponding to the intermediate productivity hypothesis. Competition represented by community mean Grime C showed a negative correlation with plant richness. Besides position, organic carbon was the most important variable for dark diversity, indicating that in late succession habitats such as forests and shrubs, dark diversity is generally low. The importance of Grime C indicate that intermediate disturbance, such as grazing, may facilitate higher species richness and lower dark diversity. Comparing various biodiversity metrics and their influencing factors might reveal important drivers of biodiversity changes and result in better conservation decision-making.

## Introduction

The global biodiversity crisis represents one of the most critical challenges in the 21^st^ century, with up to one million plant and animal species facing extinction and accelerating declines despite numerous international agreements and management responses (Tittensor et al., 2014, Butchart et al., 2010, Díaz et al., 2019). Achieving conservation goals and prioritizing efforts requires appropriate metrics to quantify biodiversity and identify the factors driving the declines. The most commonly used measure is observed species richness which traditionally depends on visual surveys to count the individual species. Although observed diversity can provide valuable insights into the richness of species within a given site, it does not account for the absent part of the species pool that could potentially inhabit that site based on suitable environmental conditions and biogeographic history, i.e., the dark diversity (Pärtel et al., 2011). Identifying this part of the biodiversity can provide a more thorough understanding of the processes influencing biodiversity and help evaluate the restoration potential of local habitats (Lewis et al., 2016).

In contrast to observed diversity, dark diversity focuses on the portion of diversity potentially able to occur in a particular habitat type. This metric can provide insight into the determinants of missing species by helping us understand why certain species are missing more often than others and the characteristics of sites typically missing many species that could potentially exist there. Quantifying dark diversity patterns, in combination with observed diversity patterns, may allow researchers to better understand the mechanisms and processes acting on individual populations or entire communities (Pärtel et al., 2017b). So far, the potential value of dark diversity to guide conservation and restoration planning has been demonstrated for mammals (Estrada et al., 2018), sharks (Boussarie et al., 2018), and fungi (Pärtel et al., 2017b, Pärtel et al., 2017a). However, most of its potential as a conservation tool has been realized in plants and has been used in conservation prioritization of plant communities across Europe (Moeslund et al., 2017) and determining the relationship between dark diversity and the invasion potential of alien species in semi-natural grasslands (Bennett et al., 2016). Dark diversity has also proven valuable in understanding plant diversity patterns, such as determining that vascular plant dark diversity across Europe follows a latitudinal gradient (Ronk et al., 2015). So far, the plant traits likely to increase a species’ probability of being part of the dark diversity include stress intolerance, tall, adaption to low light and nutrient levels, and producing fewer and heavier seeds (Riibak et al., 2017, Moeslund et al., 2017, Riibak et al., 2015). Understanding the ecological processes governing plant dark diversity is important since vascular plants can not only predict biodiversity across environmental gradients and broad taxonomic realms, but are also related to the occurrence of regionally red-listed species of other taxa (Brunbjerg et al., 2018). Furthermore, plants are bio-indicators of their abiotic environment and anthropogenic impact (Bartelheimer and Poschlod, 2016), and they form the living and dead organic carbon pools and biotic surfaces that are the niche space for all other taxonomic groups (Brunbjerg et al., 2017b). Nevertheless, as a relatively new concept, more research is required to establish its full potential and to understand the ecological processes governing dark diversity across plant communities.

Dark diversity can also be used to derive community completeness, a relativized biodiversity index, which has been proposed as a valuable tool for facilitating biodiversity comparisons irrespective of regions, ecosystems, and taxonomic groups (Pärtel et al., 2013). The community completeness index can be defined, in general terms, as the proportion of species from the regional species pool which have dispersed to and established at a site after abiotic and biotic filtering (Pärtel et al., 2013). Since patterns in observed species richness may mimic patterns in dark diversity (e.g., exhibit a strong latitudinal gradient) (Aning, 2017, Ronk et al., 2015, Pärtel et al., 2011, Zobel, 1997), community completeness can provide a better measure of biodiversity as it accounts for the variation in species pool size and expresses biodiversity on a relative scale (Pärtel et al., 2013). For instance, in previous studies, completeness exhibited no relationships to latitudinal gradients, but strong relations to anthropogenic disturbance (higher completeness in areas with lower disturbance) for fungi (Pärtel et al., 2017b), plants (Ronk et al., 2015, Ronk et al., 2016), and birds (Cam et al., 2000). Comparing the environmental processes influencing these biodiversity measurements can provide valuable information for better prioritization of resources and understanding patterns of biodiversity. However, despite observed species richness and its determining factors being relatively established, dark diversity and its completeness counterpart are new methodologies, and as such, have not been well investigated and the factors influencing them are not fully understood.

Since biodiversity varies greatly across ecosystems and are highly dependent on the habitat and region of interest, dark diversity and completeness aims to reconcile the role of local (biotic interactions, abiotic filters, dispersal, stochastic events) and large-scale processes (species diversification and historic migration patterns) underlying biodiversity patterns (Pärtel, 2014, Pärtel et al., 2011). Determining the set of species which can theoretically inhabit a site, the species pool, is typically estimated using species co-occurrence patterns with Beal’s smoothing which assumes species with shared ecological requirements and biogeographical history will have similar likelihoods of being present at a particular site (de Bello et al., 2012, Lewis et al., 2016, Beals, 1984, Münzbergová and Herben, 2004, McCune, 1994). This approach also assumes to account for prevalent competitive interaction which is a major factor influencing species occurrence patterns (de Bello et al., 2012, Cornell and Harrison, 2014), especially in plant communities (Riibak et al., 2015). Despite co-occurrence being successfully implemented as a proxy for species ecological requirements and competition, this assumption has not been examined. Another potential issue is that many ecological processes are scale dependent with different spatial scales inherently including varying amounts of environmental heterogeneity (Scott et al., 2011). However, most of dark diversity research has ignored the variability between types of habitats and have mostly been restricted to narrow sets of variables and specific habitats (Riibak et al., 2015) or largescale landscapes (Ronk et al., 2016, Ronk et al., 2015), with no studies examining how dark diversity varies across large environmental gradients or the importance of local conditions and biotic interactions. One way to consider the roles these factors play in dark diversity measurements can be provided with the recently developed ecospace framework which recognizes the influence of environmental gradients (defined as position), availability of biotic resources (defined as expansion), and the spatiotemporal extent of habitats (defined as continuity) in determining biodiversity (Brunbjerg et al., 2017b). This framework can help us better quantify and determine the different aspects that each contribute and how they relate to the various diversity metrics.

In this study, we investigate the importance of local environmental conditions that determine dark diversity and completeness in plant communities across broad environmental gradients and compare these measures with the observed richness. We use the ecospace concept to investigate how abiotic condition, biotic resources and spatiotemporal processes relate to these biodiversity measures. To examine the assumption that co-occurrence can be a proxy for competitive exclusion, we used the community mean of plant competitive scores (Grime, 1979) to quantify the importance of local interspecific competition for establishment of species. We discuss how dark diversity can contribute with new aspects for informed conservation and management.

## Materials and Methods

### Study sites

Our data stems from Biowide, a nationwide survey of biodiversity in Denmark (Brunbjerg et al., 2017a). A total of 130 study sites (40 m × 40 m) were evenly distributed across five geographic regions in Denmark with a minimum distance of 500 m between sites (Fig 1a). Each site is sampled in four 5 m circle plots (Fig 1b). Sampling was designed with the purpose of evaluating the ecospace framework, stating that biodiversity varies according to abiotic conditions, build-up and diversification of organic resources and spatio-temporal continuity (Brunbjerg et al., 2017b). The sites were stratified according to environmental gradients and 30 sites were allocated to represent cultivated habitats and 100 sites to natural and semi-natural habitats. The cultivated subset was stratified according to major land-use types and the natural subset was stratified according to soil fertility, soil moisture and successional stage. Saline and fully aquatic habitats were deliberately excluded, but temporarily inundated depressions, as well as wet mires and fens were included. The final set of 24 environmental strata consisted of the following six cultivated habitat types: Three types of fields (rotational, grass leys, set aside) and three types of plantations (beech, oak, spruce). The remaining 18 strata were natural and semi-natural habitats, constituting all factorial combinations of: fertile and infertile; dry, moist, and wet; open, tall herb/scrub, and forest. All 24 strata were replicated in each of the five geographical regions. For the purpose of the present study, we exclude all agricultural fields, resulting in a dataset of 115 sites. All field work and sampling was conducted in accordance with Responsible Research at Aarhus University and Danish law. For a thorough description of site selection and stratification procedures, see Brunbjerg et al. (2017a).

**Figure 1.**
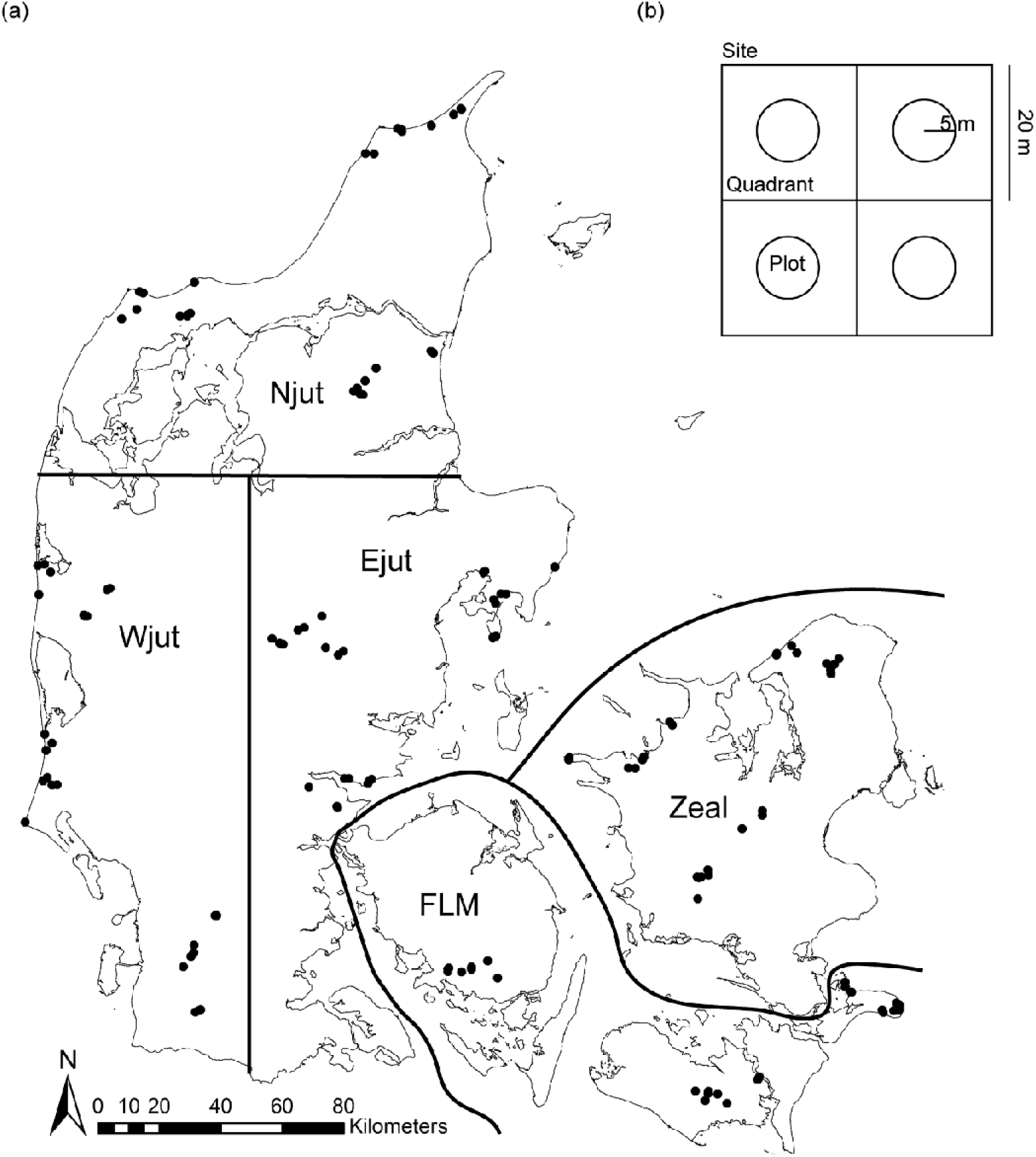
(a) Map of Denmark showing the 130 surveyed sites and the regions. (b) 40 m × 40 m site with the four quadrants and 5m circle plots.

### Data

#### Plant species richness

Vascular plant species richness was inventoried by trained botanists during the summer 2014 and spring 2015 to account for variations in phenology. The reference dataset of plant inventories in 5 m circles from the national monitoring program (Danish Nature Agency, 2016) was also inventoried by trained botanists. We removed all subspecies, hybrids, variations, and neophytes (i.e. species that are not considered a natural part of the vegetation given their history and dispersal ability, see appendix tables 6–8 in(Buchwald et al., 2013). Plant species nomenclature was obtained from the species checklist Denmark database from https://allearter-databasen.dk to match the two datasets and account for different synonyms.

#### Explanatory variables

We used the ecospace concept to investigate how abiotic condition (i.e., position), biotic resources (i.e., expansion), and spatiotemporal extent of habitats (i.e., continuity) explain plant richness, dark diversity and community completeness (Table 1). The position variables included in the model were soil moisture index (SMI) and soil fertility index (SFI). For each site SFI represents the predicted value from the best linear model (of all sites) of site mean Ellenberg N (plant-based bioindication of nutrient status; (Ellenberg et al., 1991)) as a function of soil Ca, leaf N, leaf NP and soil type. We calculated a soil moisture index for each site using the predicted values from the best linear model (of all sites) of mean Ellenberg F (plant-based bioindication of soil moisture; Ellenberg et al., 1991) as a function of mean precipitation in 2001-2010 (10 km × 10 km grid resolution) and measured site soil moisture (trimmed mean of 16 measures pr. site taken with a FieldScout TDR 300 Soil Moisture Meter in May 2016) (Brunbjerg et al., in prep). Position also included soil pH measured on 4 pooled soil samples from 0-10 cm depth and light measured as light intensity (Lux) using HOBO Pendant^®^ Temperature/Light 8K Data Loggers installed at the ground as detailed in Brunbjerg et al. (2017a). The expansion variables included: 1) Bare soil percent coverage as a subjective estimate, 2) Litter mass (g/m-2 of four litter samples within a 21 cm × 21 cm frame pr. site). The four samples from each site were pooled, taken to the laboratory, dried (60° for 48 hours) and mass (g/m-2) was registered, 3) Soil organic matter as a percentage of the 0-10 cm soil core that was categorized as organic soil, and 4) soil organic carbon as % soil C in 0-10 cm soil layer (g/m-2 average of four soil samples taken in each site) as described in Brunbjerg et al. (2017a). Expansion also included structural heterogeneity from variation in the digital elevation model (DEM6) (40 cm × 40 cm resolution) from the digital surface model (40 cm × 40 cm resolution) to create a grid representing the above-ground vegetation height. From this the structural heterogeneity was calculated as the sum of variability in the shrub layer and canopy layer, measured as the variance of the 90th percentile for returns > 3 m within the site reflecting the variability of the height of mainly trees and the variance of the 90th percentile for returns 30 cm - 3 m reflecting the variability of the height of the shrub layer (Brunbjerg et al., in prep). Landscape characteristics were represented by two co-variables: 1) share of intensive fields within a 500 m buffer and 2) the share of natural habitats in a 1 km × 1 km grid from a national mapping, interpolated using Spline in ArcGIS 10.2.2 (weight 0.5, number of points 9, (Ejrnæs et al., 2014)). Temporal continuity was estimated as time since major land use change within the 40 m × 40 m site. For each site, a temporal sequence of aerial photos and historical maps was inspected starting with the most recent photos (photos from 2014, 2012, 2010, 2008, 2006, 2004, 2002, 1999, 1995, 1968, 1954, 1945) and ending with historical maps reflecting land use in the period 1842-1945. Temporal continuity (the year in which a change could be identified) was reclassified into a numeric 4-level variable: 1: 1-14 years, 2: 15-44 years, 3: 45-135 years, 4: >135 years (Brunbjerg et al., 2017a). Lastly, to examine the importance of species interactions and the assumption that co-occurrence can be a proxy for competitive exclusion, we quantified local interspecific competition by calculating an index for the intensity of plant competition (Grime C) using the community mean plant competitive score (Grime, 1979) which is thought to reflect plant species’ adaptation to interspecific competition (Grime et al., 2014). The original C-S-R species strategies were recoded into numeric mean site C values (Ejrnæs and Bruun, 2000).

**Table 1.** Explanatory variables, their affiliation to ecospace elements and their hypothesized association with response variables. Question marks indicate that hypotheses are either ambiguous or unknown.

| Explanatory variable | Ecospace dimension | Dark Diversity | Plant Richness |
| --- | --- | --- | --- |
| Soil moisture index (SMI) | Position | ? | Unimodal |
| Soil fertility index (SFI) | Position | ? | Unimodal |
| Soil pH | Position | ? | Positive |
| Light | Position | ? | Positive |
| Litter | Expansion | Positive | Negative |
| Bare soil | Expansion | Negative | Positive |
| Organic carbon (OrgC) | Expansion | ? | ? |
| Organic matter | Expansion | ? | ? |
| Shrub height variation | Expansion | Negative | Positive |
| Canopy height variation | Expansion | Negative | Positive |
| Intensive field | Continuity | Positive | Negative |
| Natural landscape | Continuity | Negative | Positive |
| Temporal continuity | Continuity | Negative | Positive |
| Competition (Grime C) | Interactions | Positive | Negative |

### Data analysis

#### Regional pool, dark diversity and completeness

All statistical analyses were performed in R version 3.5.3 (R Core Team, 2019). To calculate regional pools, we used 5 m circle plots of observed plant species from both datasets. We did not include species-poor plots, i.e., those with less than five observed species, i.e., resulting in 448 plots from Biowide and 52362 plots from the reference data set. The regional pool was calculated using Beals index (Beals, 1984), as recommended by (Lewis et al., 2016). The Beals index represents the probability that a particular species will occur within a given site based on the assemblage of co-occurring species (Beals, 1984, Münzbergová and Herben, 2004, McCune, 1994). We calculated Beals index using the ‘beals’ function in the ‘vegan’ package (Oksanen et al., 2017). The threshold for including a particular species in the regional species pool is recommended to be the 5^th^ percentile of the Beals index value for the species in question (Gijbels et al., 2012, Ronk et al., 2015). Preceding the calculation of each threshold, the lowest Beals index value among plots with occurrence of the species in question was identified, and all plots having values below that minimum were not considered.

Analyses were done at the site level (n=115) by creating a site regional pool combining the four plot regional pools at each site. Observed species in the site, but not included in the regional pools (n=2) were added to the regional pools to ensure that site regional pool included all observed species. Then, dark diversity was calculated for each plot as the difference between the regional pool and the observed species richness (Pärtel et al., 2011) and completeness was calculated following Pärtel et al. (2013) using the formula *ln(observed richness/dark diversity)*. Because dark diversity is relative based on habitat types and may not be suitable for comparison across habitats (Scott et al., 2011), completeness is suggested as possible alternative (Pärtel et al., 2013). Completeness should have been suitable for investigations across habitat types, but in this study we found that completeness was highly correlated with observed species richness (Figure 2). Therefore, we obtained a dark diversity measure corrected for habitat differences by using the residuals of a model of dark diversity as a function of the habitat type, which we termed Bio-stratum, a classification of the sites using all inventoried species data, i.e., plants, fungi, insects, etc., resulting in eight classes spanning gradients in succession (Early, Late), moisture (Wet, Dry) and nutrients (Rich, Poor). The habitat types explained 51% of the variation in dark diversity.

**Figure 2.**
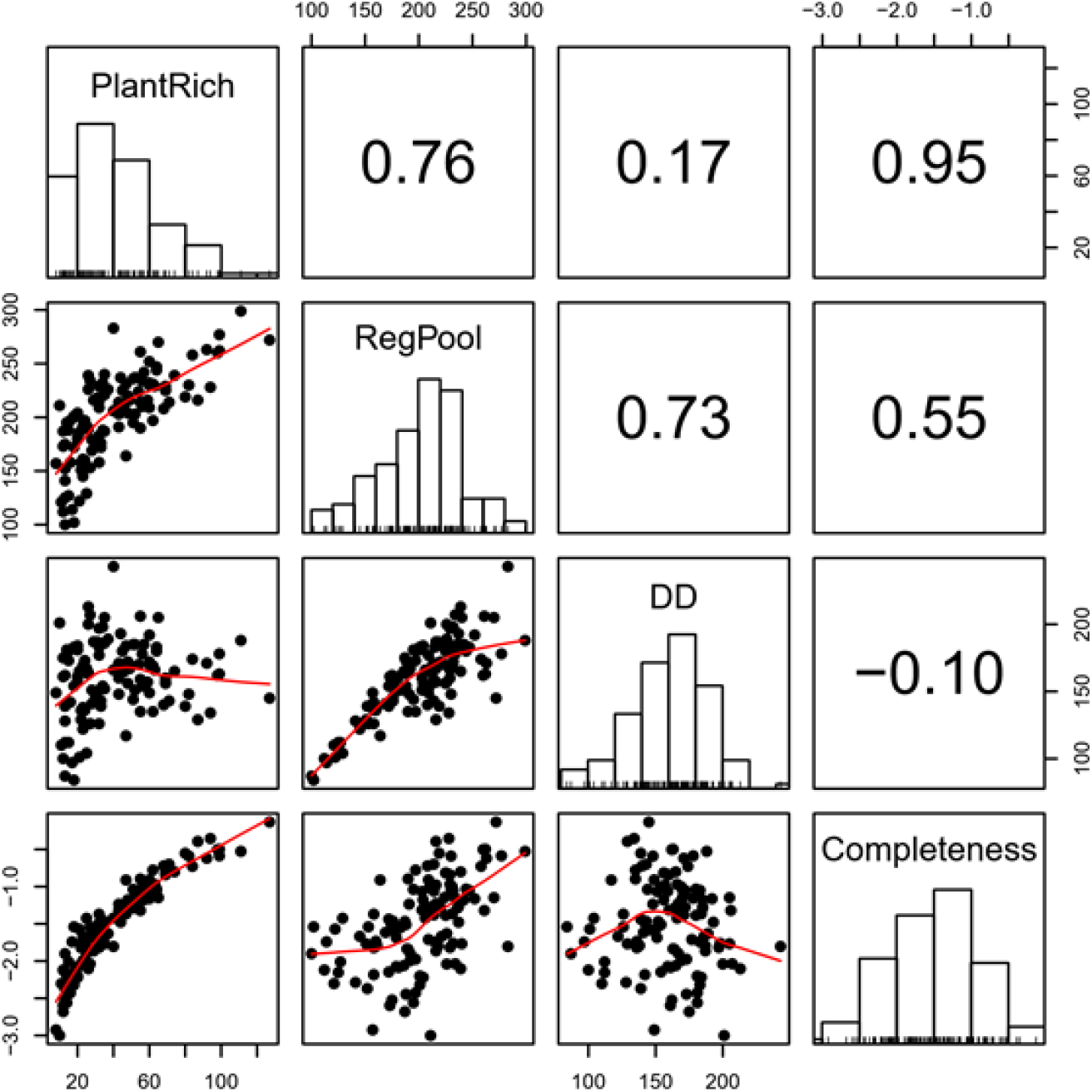
Spearman rank correlations and dot plots of site regional pool, plant species richness, dark diversity and completeness. The red line in the plot shows a loess smoothing line.

#### Statistical analyses

Soil pH, litter mass, organic carbon, organic matter and shrub and canopy height variation were log-transformed and all explanatory variables were standardized. We conducted model selection by first testing for multicollinearity, which could affect p-values and model validity using the Variance Inflation Factor (VIF). We then removed canopy height variation and organic matter resulting in a maximum VIF of 2.9. The remaining variables were used as explanatory variables (linear terms) in a set of generalized linear models (GLM) using the MASS package (Venables and Ripley, 2013) with Poisson distribution for the count data for dark diversity and plant richness, and normal distribution for residual dark diversity. However, since models for plant richness were over-dispersed we chose negative binomial models for this response instead. To allow for non-linear relationships for position variables corresponding to the intermediate disturbance hypothesis (Townsend et al., 1997, Connell, 1978) and intermediate productivity hypothesis(Fraser et al., 2015), we used AIC (Burnham and Anderson, 2002) to evaluate if inclusion of quadratic terms for the variables SMI, SFI, Light, soil pH and bare soil improved the model fit, in which case relevant quadratic terms were added. Subsequently, we checked the parameter estimates against plausible ecological hypotheses (Table 1) and excluded ecologically implausible responses (Burnham and Anderson, 2002). We then dropped remaining variables sequentially based on AIC using the lme4 package (Bates et al., 2007). The final models were tested for over-dispersion and evaluated by visual inspection of residual plots and for spatial autocorrelation using Moran’s I in the ape package (Paradis et al., 2004).

## Results

We found a total of 580 species of vascular plants in the 115 sites spanning from open habitats to shrubs and late succession forests. The species richness per site ranged from 8 to 127 species and dark diversity ranged from 84 to 243 species. Completeness and plant species richness were highly positively correlated, whereas dark diversity was less correlated with plant species richness (Figure 2). The final models explained between 14 and 65 % of the variation in dark diversity, residual dark diversity and species richness (Table 2). We found position variables to be important for dark diversity and plant species richness (Figure 3, 5). SMI invoked a unimodal response in dark diversity but a bimodal response in species richness. We observed a positive effect of SFI and soil pH on dark diversity and unimodal relationships with species richness. Light had a negative linear and unimodal relationship with dark diversity and species richness, respectively. No position variables were found to be important for the residual dark diversity, as would be expected. Organic carbon was important for all measured responses with a linear negative relationship across response variables (Figure 3-5). Grime C had a positive linear relationship with dark diversity and a negative linear relationship with plant species richness (Figure 3, 5), while natural landscapes had a linear negative relationship with residual dark diversity (Figure 4).

**Table 2.** Regression results for dark diversity (DD) using Poisson, residual dark diversity for habitat types using ordinary least squares and plant species richness (PlantRich) with negative binomial. R^2^ is calculated as 1-(model deviance/model null deviance) for dark diversity and plant species richness for residual dark diversity (DD) we report the multiple R^2^. Parentheses gives the standard errors. *p<0.1; **p<0.05; ***p<0.01.

| Ecospace | Variables | DD | Residual DD | Plant richness |
| --- | --- | --- | --- | --- |
|  | Intercept | 5.090<sup>***</sup> (0.014) | -0.000 (0.136) | 3.762<sup>***</sup> (0.081) |
| Position | Soil moisture index (SMI) | -0.020<sup>*</sup> (0.011) | n.s. | 0.190<sup>***</sup> (0.054) |
| Position | SMI<sup>2</sup> | -0.025<sup>**</sup> (0.012) | n.s. | 0.150<sup>***</sup> (0.053) |
| Position | Soil fertility index (SFI) | 0.069<sup>***</sup> (0.011) | n.s. | 0.233<sup>***</sup> (0.055) |
| Position | SFI<sup>2</sup> | n.s. | n.s. | -0.145<sup>***</sup> (0.029) |
| Position | Soil pH | 0.030<sup>***</sup> (0.009) | n.s. | 0.269<sup>***</sup> (0.071) |
| Position | Soil pH<sup>2</sup> | n.s. | n.s. | -0.060<sup>*</sup> (0.034) |
| Position | Light | -0.042<sup>***</sup> (0.010) | n.s. | 0.102 (0.066) |
| Position | Light<sup>2</sup> | n.s. | n.s. | -0.086<sup>*</sup> (0.047) |
| Expansion | Litter (log) | n.s. | n.s. | -0.105<sup>*</sup> (0.056) |
| Expansion | Organic carbon (OrgC) | -0.045<sup>***</sup> (0.011) | -0.465<sup>***</sup> (0.137) | -0.126<sup>**</sup> (0.050) |
| Expansion | Bare soil | n.s. | n.s. | -0.079 (0.050) |
| Expansion | Shrub height variation | n.s. | n.s. | 0.161<sup>***</sup> (0.048) |
| Interaction | Competition (Grime C) | 0.037<sup>***</sup> (0.009) | n.s. | -0.192<sup>***</sup> (0.048) |
| Continuity | Natural Landscape | n.s. | -0.326<sup>**</sup> (0.137) | n.s. |
| Multiple $R^2$ | | 0.65 | 0.14 | 0.65 |

**Figure 3.**
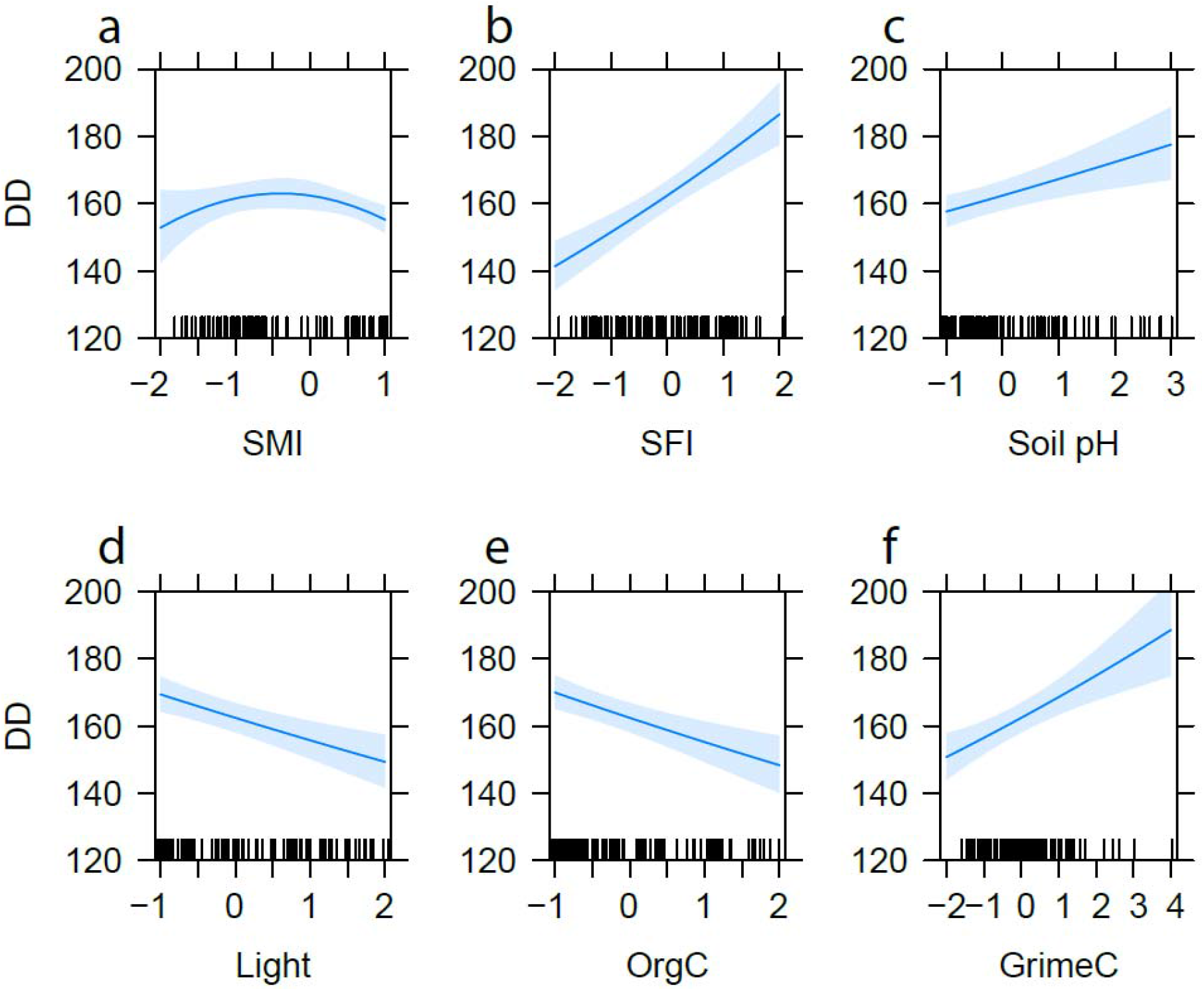
Parameter estimates with 95% confidence intervals from the significant environmental variables predicting overall dark diversity. Relationships between the dark diversity and (a) soil moisture index (SMI), (b) soil fertility index, (c) soil pH, (d) light, (e) organic matter, and (f) competition (GrimeC). Y-axis is truncated.

**Figure 4.**
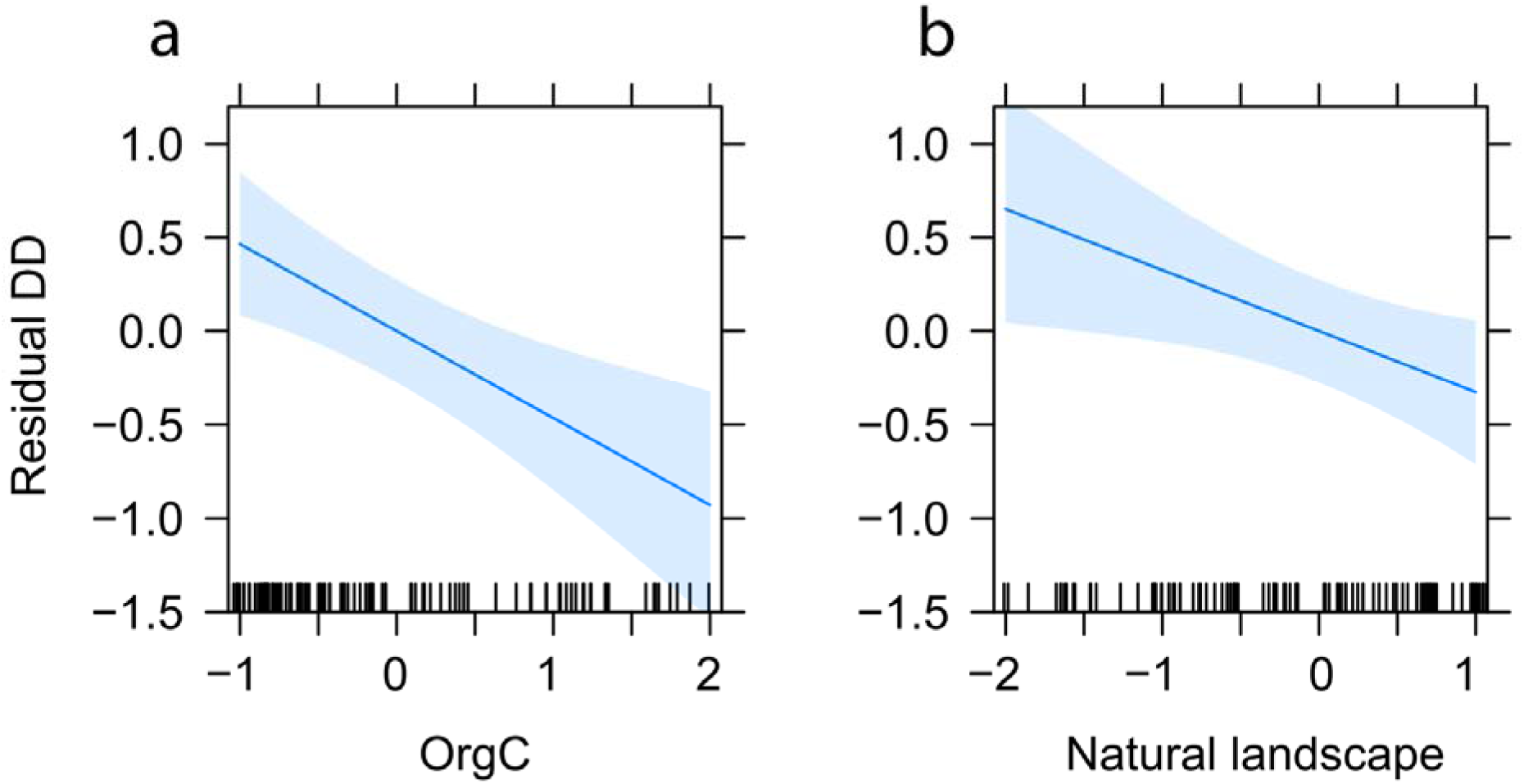
Parameter estimates with 95% confidence intervals from the significant environmental variables predicting residual dark diversity for habitat types. Relationships between residual dark diversity and (a) organic carbon (OrgC), and (b) natural landscape.

**Figure 5.**
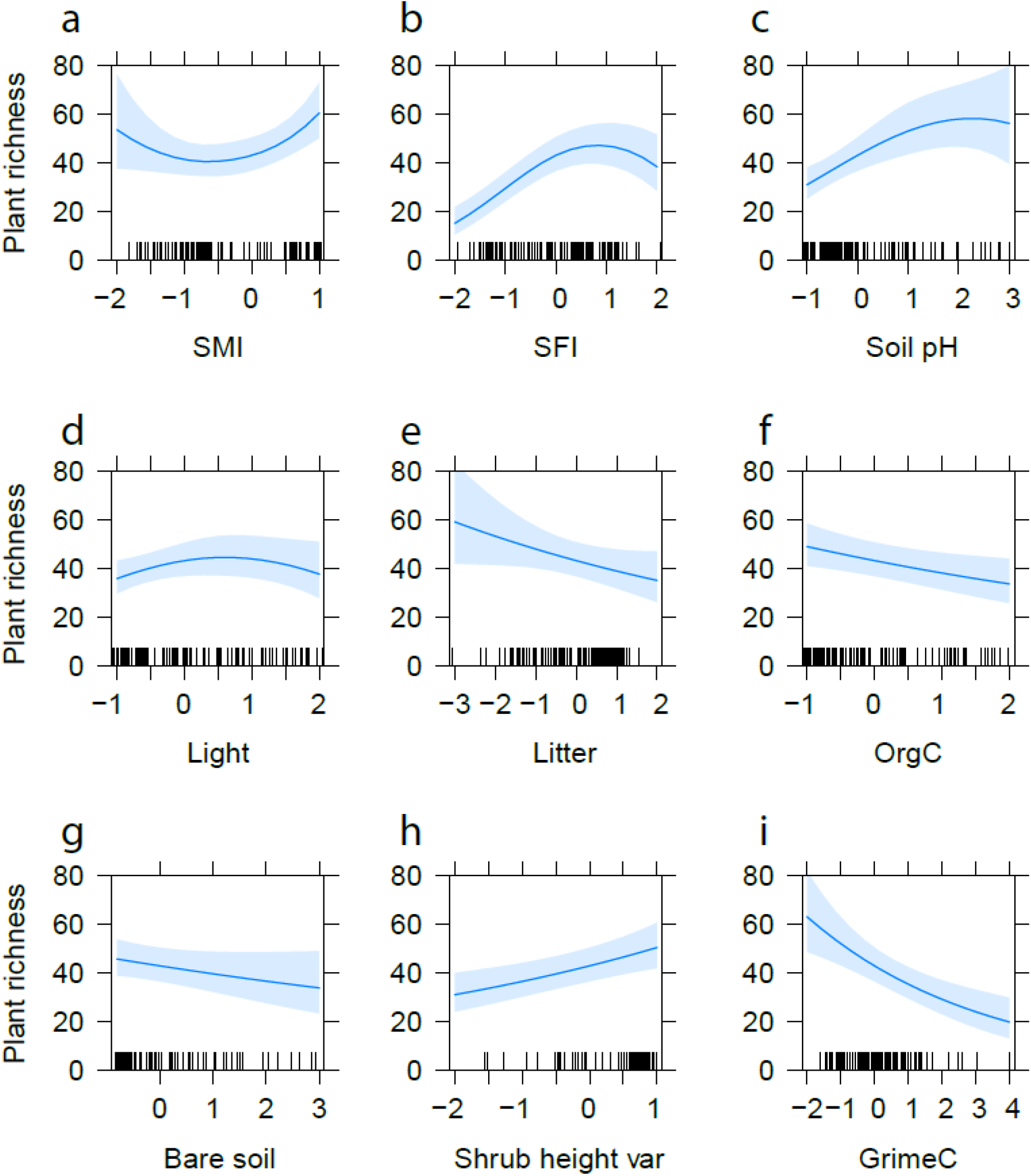
Parameter estimates with 95% confidence intervals from the significant environmental variables predicting plant species richness. Relationships between the plant species richness and (a) soil moisture index (SMI), (b) soil fertility index (SFI) (c) soil pH, (d) litter mass, (f) organic carbon, (g) bare soil, (h) shrub height variation, and (i) competition (GrimeC).

## Discussion

Completeness, thought to be less dependent on habitat types (Pärtel et al., 2013), was highly correlated with plant species richness and therefore added no new information to our study of plant community diversity aspects. Therefore, we explore residual dark diversity, another measure independent of habitat type. Position variables were important for both plant richness and dark diversity. Not surprisingly, once differences between habitat types were accounted for, i.e. residual dark diversity, the position variables were no longer significant. Plant richness was highest at intermediate conditions of soil fertility and pH, corresponding to the intermediate productivity hypothesis, which states that few species can tolerate the environmental stresses at low productivity and a few highly competitive species dominate at high productivity (Fraser et al., 2015). Species richness increased with pH, possibly corresponding to the generally large regional species pool in calcareous habitats, and aligns with previous research indicating that plant diversity has a strong positive association with soil pH in temperate and boreal regions (Pärtel, 2002, Pärtel et al., 2004). The unimodal relationship between dark diversity and soil moisture may be due to communities at the extremes being more distinct than communities at intermediate soil moisture, i.e., specific adaptations for waterlogged and very dry soil are required (Ernst, 1990). Therefore, fewer coincidental species may appear in these extreme habitats compared to habitats of intermediate moisture, resulting in a low estimated regional pool and lower dark diversity at the extremes.

Regarding the expansion variables, variation in shrub height was negatively correlated with dark diversity and positively correlated with species richness, indicating that vegetation heterogeneity (and resulting local variation in light, temperature, etc.) increases the establishment and survival of species (Stein et al., 2014). Previous research has shown that shrub height can increase species richness in competitive environments as variance in heights can ameliorate conditions to neighboring plants and cause biomass to be distributed in vertical space, thereby reducing competition for space (Bråthen and Lortie, 2016). We also found bare soil had a negative effect on species richness as would be expected since more bare soil will by definition have fewer observed individuals and species. The only expansion variable to influence both dark diversity measures (and the only expansion variable influencing the residual dark diversity) and species richness was soil organic carbon, which had a negative relationship with both dark diversity and species richness. Previous studies have also indicated that native and exotic richness is lower with higher soil organic carbons (Perelman et al., 2007). These results may be due to temporal continuity and the effect on the various successional levels on the accumulation of soil organic carbon. For example, more established and complete communities (e.g., old forests) accumulate greater organic carbon but exhibit lower species diversity (Garnier et al., 2004). This may also be true for species poor acidic habitats like mires, bogs and heaths, which were represented in this study. Competition plays a large role with early successional species able to modify the environment through rapid growth and inhibit the success of species currently present, while later stage communities are typically composed of the most competitive species and replace earlier succession species (Connell and Slatyer, 1977). Therefore, the potential species that can inhabit the area may be restricted by competitive species. This competitive advantage inherit in late successional communities results in greater dark diversity, corresponding to our result that Grime C is important for plant dark diversity. The positive effect of Grime C on dark diversity indicates that there are more species missing from communities dominated by competitive species. The effect on plant species richness was opposite, likely because dominant competitive species exclude other species, thereby reducing the overall plant species richness. It seems that competition does not affect the species pool, otherwise suitable species would become locally extinct due to dominant species, but they still remain in dark diversity and can be restored if competition is controlled (e.g. disturbance). Spatial continuity, in this case natural landscape, was only important for residual dark diversity, whereas competition was not important. One possibility is that the habitat types account not only for variation in position but also the inherent competitive strategies of the species in the habitat types e.g., late successional stages dominated by competitive species, possibly explaining why competition is not important for residual dark diversity. Natural landscapes may decrease dark diversity by influencing local processes, i.e., landscapes with high nature density are likely to have a higher local pool of species, increased dispersal, increased species survival, metapopulation structures, and less negative edge-effects of intense land use (Brunbjerg et al., 2017b). Another explanation why natural landscape was only significant for residual dark diversity may be that the effect of habitat type and position variables it is large and masks all other effects in both dark and observed diversity. This effect is only visible when these variables are accounted for.

This study shows that there are many different methodologies to measure biodiversity, and that they contribute with different aspects to better understand drivers of diversity, with their applicability depending on the desired objectives and goals. For example, if looking at the effects of organic carbon on species richness one may conclude that carbon capture and storage may lead to loss of species richness, however dark diversity indicates that carbon storage could increase habitat completeness. Just like there are dozens of different ways to measure species diversity (Lyashevska and Farnsworth, 2012), there are also many ways to calculate dark diversity. Besides the typical co-occurrence measure, dark diversity can also be calculated as community completeness (Pärtel et al., 2013), based on ecological requirements (Lewis et al., 2016, Bello et al., 2016) or species distribution (Bello et al., 2016), and probabilistic measurements such as hypergeometric distributions (Carmona et al., 2019). Here, we compared dark diversity and completeness, but found that completeness was highly correlated with observed species richness. We therefore advocate to compare and analyze different diversity measures as both dark diversity and completeness are still relatively new metrics. Furthermore, applying dark diversity within one habitat type, e.g., as seen for grasslands (Riibak et al., 2015) may produce adequate results, however, when applying dark diversity across habitat types or broad environmental gradients, correcting for these differences through a residual dark diversity measure may provide more interpretable results.

With global biodiversity rapidly disappearing, it is vital to understand the drivers of biodiversity to prioritize conservation and make management more efficient. In this study, besides the ecospace position variables, competition seems to be the greatest driver of plant richness. Conservation management focusing on intermediate disturbance such as grazing can disrupt competitive communities making room for more species. Besides ecospace position, organic carbon was the most important variable for both dark diversity measures indicating that advanced succession and possibly temporal continuity may increase completeness or decrease dark diversity. Examining the influencing factors of different measures of biodiversity can lead to better decision-making in the future conservation of the world’s biodiversity.

## Acknowledgements

We sincerely thank Aage V. Jensen Nature Fund for financial support to CF, AKB, JM, LD, KC and JV for the project “Dark Diversity in nature management”. The biowide project and REJ was supported by a grant from the Villum Foundation (VKR-023343). MP has been supported by the Estonian Ministry of Education and Research (IUT20–29), and the European Regional Development Fund (Centre of Excellence EcolChange).The authors declare no conflict of interest.

